# *Toxoplasma gondii* infection of neurons alters the production and content of extracellular vesicles altering astrocyte phenotype and contributing to the loss of GLT-1 in the infected brain

**DOI:** 10.1101/2024.11.08.622601

**Authors:** Emily Z. Tabaie, Ziting Gao, Stacey Gomez, Kristina Bergersen, Wenwan Zhong, Emma H. Wilson

## Abstract

*Toxoplasma gondii (T. gondii)* cyst formation in the central nervous system only occurs in neurons allowing the parasite to remain latent for the lifetime of the host. Astrocytes are fundamental to neuronal health by providing nutrients and structural support and help regulate neurotransmitters by continuous communication with neurons. It is not yet known how infection and the presence of intracellular cysts, disrupts the crucial relationship between these cells. Extracellular vesicles (EVs) function in intracellular communication and can contain proteins, lipids, DNA, miRNA, and other RNA subtypes. EVs are produced by all cells including neurons and play an important role in neuronal-astrocyte interactions including the regulation of glutamate receptors on astrocytes. Previous work has demonstrated Toxoplasma infection reduces astrocytic expression of the primary glutamate transporter, GLT-1. Here we tested if cyst infection of neurons alters the production and content of EVs. EVs were isolated from uninfected and infected primary murine cortical neurons and their size, concentration, and characterization were confirmed with nanoparticle tracking analysis (NTA), transmission electron microscopy (TEM), CD63 ELISA, liquid chromatography (LC)-mass spectrometry (MS)/MS, and microRNA Sequencing. Analysis reveals that infection of neurons reduced neuronal production of EVs and altered their protein and miRNA content. EVs from infected neurons contained secreted Toxoplasma proteins GRA1, GRA2, GRA7, MAG1 and MAG2 associated with cyst formation. Following incubation of neuronal EVs with primary astrocytes, a proportion of EVs colocalize to the nucleus. EVs from infected neurons altered gene expression of astrocytes leading to a downregulation of GLT-1 protein expression and an increase in pro-inflammatory transcriptional signatures. These results demonstrate the ability of a parasitic infection in the brain to alter EV production and the fundamental communication between neurons and astrocytes.

## 1 Introduction

Nearly one-third of the world’s population is infected with the obligate intracellular parasite, *Toxoplasma gondii (T. gondii)* (Bando et al., 2021; el-On & Peiser, 2003). Following a period of rapid replication and systemic inflammation, infection leads to slow growing tissue cysts in the brain, skeletal muscle, and cardiac muscle of the host (Mendez & Koshy, 2017). Along with the transition to the less cell lytic cyst form, chronic infection is mediated by the ability of the parasite to manipulate cell signaling pathways and inhibit host cell defense mechanisms, while remaining dormant and undetected in the infected cell.

In the brain, cysts reside intracellularly within neurons. Neurons are responsible for receiving and sending sensory and motor signals, allowing for information to be transmitted across great distances (Farhy-Tselnicker & Allen, 2018). Although Toxoplasma infection does not cause significant clinical pathology in the immune competent individual, there is evidence cyst infection of neurons are functionally compromised with a reduction in dendritic spines and electrical activity (David et al., 2016; Mendez & Koshy, 2017). The function of neurons is intimately linked with astrocytes, glial central nervous system (CNS) resident cells. Astrocytes are the most abundant glial cell in the brain and assist with support and nourishment of neurons, promoting signaling, and taking-up nutrients from the blood to transfer to neurons (Zang et al., 2022). In addition, astrocytes are vital for maintaining the blood brain barrier (BBB) and their dysfunction can lead to neurological disorders, including neurodegenerative diseases, stroke, and epilepsy (Rama Rao & Kielian, 2015). Importantly, astrocyte-to-neuron communication and function is mediated through direct cell-to-cell contact or the production, secretion and uptake of extracellular vesicles (EVs). Neurons can communicate information to other cells through electrical and chemical synapses, however, recently their ability to transfer material in EVs has established an important additional paracrine cell-cell mechanism. The communication between neurons and astrocytes is vital for various functions, including synapse formation, ion homeostasis, and the regulation of important neurotransmitters in the brain (Farizatto & Baldwin, 2023).

Glutamate is the primary excitatory neurotransmitter in the brain. Astrocytic glutamate transporter, GLT-1, is responsible for 90% of extracellular glutamate uptake in the CNS, a crucial function to prevent glutamate buildup in the synaptic cleft (Pajarillo et al., 2019). GLT-1 dysregulation results in excess glutamate concentrations resulting in an excitotoxic state that can lead to spontaneous seizure activity (Bjørnsen et al., 2007; Zhou & Danbolt, 2014). Astrocytes regulate CNS glutamate by adjusting uptake, release, and synthesis into glutamine. We have previously demonstrated that following Toxoplasma infection, there is a decrease in GLT-1 expression resulting in an increase in extracellular glutamate concentrations (David et al., 2016). GLT-1 expression is dysregulated in many neurological diseases, making its downregulation an important area of research. The exact mechanism of GLT-1 regulation has yet to be identified, however post-translational modifications and extracellular vesicles are at least partially responsible for its regulation (Men et al., 2019; Morel et al., 2013; Peterson & Binder, 2019).

EVs are membrane bound molecules that contain proteins, lipids, DNA, miRNA, and many other RNA subtypes. All cells are able to secrete EVs and they can be observed in many biological fluids including blood, tear, saliva, urine, breast milk, and cerebrospinal fluid (Gioseffi et al., 2021; Marcilla et al., 2014). Parasites, including *T. gondii*, also have the capability of producing their own EVs (Maia et al., 2021; Quiarim et al., 2021). EVs are derived from the fusion of multivesicular bodies with the plasma membrane and upon the extracellular release of the intraluminal vesicles, the EVs can dock to the plasma membrane of the target cell and enter through fusion or endocytosis mechanisms (Raposo & Stoorvogel, 2013). EVs release their content and alter signaling pathways to assist with cellular communication, through an unknown process. One main characteristic of EVs is that they can function in intercellular communication without the need for cell-to-cell contact or specific receptor interactions.

Here we describe the ability of Toxoplasma infection to alter the production and content of EVs from infected neurons and the resulting effect on recipient astrocytes. Results demonstrate a significant decrease in the concentration of EVs from *T. gondii* infected neurons and an alteration in EV protein and miRNA content. EVs from infected neurons altered astrocytes, including an upregulation in inflammatory genes and a significant decrease in astrocytic GLT-1 protein expression. These processes reveal a previously unknown mechanism of parasite-induced changes in the brain that have the potential to broadly alter neurochemistry in the infected CNS.

## 2 Materials and Methods

### 2.1 Animals

All mice were stored in accordance with the Animal Welfare Act and all efforts were made to minimize suffering. All protocols were approved by the Institutional Animal Care and Use Committee (IACUC) at the University of California, Riverside. SW, CBA, and C57BL/6 mice were obtained from Jackson Laboratories (Jackson ImmunoResearch Laboratories, Inc., West Grove, PA, USA) and maintained in a pathogen-free environment according to IACUC protocols at the University of California Riverside.

### 2.2 Cell Culture

Primary cortical neuron cultures were grown from the cortex of C57BL/6 embryos (E18-20). The forebrain was removed, and meninges stripped in Hanks’ Balanced Salt Solution (HBSS) without Ca^2+^ and Mg^2+^. The tissue was spun at 300 g for 2 minutes at room temperature. Neural tissue dissociation kit (Miltenyi Biotec, 130-093-231) was used according to the manufacturer’s protocol. Tissue was incubated with enzymes at 37°C with 5% CO_2_ under slow, continuous rocking conditions and dissociated through an 18-gauge needle followed by a 20-gauge needle followed by further incubation. The tissue was filtered through a 40 μm strainer and washed in HBSS with Ca^2+^ and Mg^2+^. Cells were spun at 300 g for 10 min at room temperature and the pellet was resuspended in neurobasal medium (supplemented with 2% B-27 supplement, 0.5 mM L-glutamine, and 1% penicillin-streptomycin). Cells were plated on poly-L-lysine (PLL) coated 6 well plates at 5.0×10^5^ cells per well in 2 mL and incubated at 37°C with 5% CO_2_ in complete neurobasal medium supplemented with 25 M of L-Glutamic Acid. Half of the medium was changed every three days with neurobasal medium and filtered through a 0.8 μm filter and stored at −80°C.

Primary cortical astrocyte cultures were grown from the cortex of C57BL/6 pups (P0-3). The forebrain was removed, and meninges stripped in an isolating medium (1X PBS with 0.1% BSA (bovine serum albumin) and 0.45% glucose). Cortices were mashed and filtered through a 40 μm strainer in washing medium (Dulbecco Minimum Essential Medium (DMEM) supplemented with 2% HI-FBS (heat-inactivated fetal bovine serum)). The cells were spun at 2,000 rpm for 10 minutes at 4°C and washed three times. The final pellet was resuspended in astrocyte culture medium (DMEM supplemented with 10% HI-FBS, 1% non-essential amino acids, 1% L-glutamine, 1% 4-(2-hydroxyethyl)-1-piperazineethanesulfonic acid buffer (HEPES), and 1% penicillin/streptomycin) and plated in T25 flasks incubated at 37°C with 5% CO_2_. Astrocyte medium was changed every 2 days until day 8. On day 8 the flasks were shaken for 2 hours at 260 rpm at 37°C. The astrocyte medium was changed, and the flasks were then shaken for 24 hours at 100 rpm at 37°C. The cells were washed with HBSS without Ca^2+^ and Mg^2+^ and then lifted with 0.25% Trypsin-EDTA (disodium ethylenediaminetetraacetic acid). The reaction was stopped with astrocyte medium, and the cells were centrifuged at 1,200 rpm for 5 minutes at room temperature. The pellet was resuspended in astrocyte medium and plated on 6 well plates at 1.0×10^5^ or 2.5×10^5^ cells/mL in 2 mL and incubated at 37°C with 5% CO_2_.

### 2.3 Infections

The Me49B7 strain of *T. gondii* was maintained in cyst form through continuous passage in SW and CBA background mice. For *in vitro* cultures the parasite was maintained in human foreskin fibroblasts (HFF) in parasite medium (DMEM supplemented with 5% HI-FBS, 1% L-glutamine, 1 mL/L gentamicin, and 1% penicillin/streptomycin) and incubated at 37°C with 5% CO_2_. Once confluent the neurons were infected with the parasite at day 8. Different MOIs (multiplicities of infection) of 0.25, 0.50, 0.75, and 1.0 were tested and a final MOI of 0.50 was chosen due to less dendrite loss as calculated by Neurolucida (**Supplemental Figure 1**). Day 9, 50% of the media was collected, filtered through a 0.8 μm filter, and stored at −80°C for further EV isolation. On Day 11, all 2 mL of the media was collected, filtered, and stored at −80°C until further isolation.

### 2.4 EV Isolation

Cyst formation occurs rapidly in neurons following infection. At 3 days post infection the supernatant was collected and EVs isolated through a series of ultracentrifugation methods and a membrane affinity ExoEasy isolation kit (Théry et al., 2018). To first remove debris, the supernatant was spun at 500 g for 15 minutes at 4°C. To next remove apoptotic bodies, the supernatant was collected and run on the ultracentrifuge at 15,000 g for 20 minutes at 4°C (Doyle & Wang, 2019; Görgens et al., 2022). Lastly, EVs were isolated using the ExoEasy Kit from Qiagen (76064), according to the manufacturer’s protocol. Mouse embryo fibroblast EVs from C57BL/6 mice were purchased from STEMON (EDA04-00) to act as a positive EV control. EV isolation and quantification was in line with MISEV guidelines (Welsh et al., 2024). Collected EVs were counted on the NanoSight NS300 (Malvern Instruments). Isolated EVs were diluted in 1 mL of 1X PBS in a 100X dilution and run on a nanoparticle tracking analysis (NTA). The infusion rate and withdrawal range were both set to 1,000. The screen gain was set to 10 and the detection threshold was set to 5, along with a focus of 60 for all recordings. The NTA provided an average size and concentration of the particles in each sample. Each experiment was performed in technical triplicates.

### 2.5 Transmission Electron Microscopy (TEM)

The presence of EVs was analyzed by TEM. Carbon-coated 200-mesh copper grids with continuous film were glow discharged (EMS 100x) for 20 seconds. To the grids 10 μL of EVs were added until dry. The grids were treated with 30 μL of 1% uranyl acetate for three different time increments. First for 1 minute, then 30 seconds, and a final 30 seconds. In between each incubation time the uranyl acetate was removed with filter paper before a fresh drop was added. The grids were dried for 10 minutes before imaging. Grids were observed under a Talos L120C TEM (Thermo Scientific) operating at 120 kV. Images were analyzed using ImageJ software.

### 2.6 ELISA

A half-area, high affinity binding 96 well plate was coated with a CD63 capture antibody (Abcam, ab253503) and incubated overnight at 4°C. The plate was then blocked with 1% BSA in 1X PBS for an hour at room temperature. Varying dilutions of sample (1:50, 1:100, and 1:200) were added, along with CD63 standards (Sino Biological, 50557-MNCH), and incubated for 1 hour at 37°C. A Biotinylated-CD63 detection antibody (Abcam, ab253503) was added and incubated for 1 hour at 37°C, protected from light. Streptavidin-HRP was added for 20 minutes, followed by a substrate solution for another 15 minutes. Stop solution was added and the plate was read at 450 nm. Samples were plated in triplicate.

### 2.7 Immunocytochemistry (ICC)

Primary murine cortical neurons were stained 11 days after plating. The supernatant was aspirated, and the cells were washed once with 1X PBS. The cells were fixed with 4% paraformaldehyde (PFA) for 15 minutes rotating, covered from light at room temperature. The solution was aspirated, and coverslips washed three times with 1X PBS. Permeabilization buffer (deionized water supplemented with 10% 10X PBS, 3% BSA, and 0.3% Triton X-100) was added for 10 minutes under rotation, covered from light at room temperature, the solution aspirated, and the coverslips washed three times with 1X PBS. Addition of 5% Donkey Serum (DS), in 1X PBS was used for blocking followed by primary antibody ((chicken MAP2 (Abcam, ab5392, 1:1,000), Biotinylated DBA (Vector Labs, B-1035, 1:500), and rabbit NeuN (Abcam, ab177487, 1:500)) diluted in blocking buffer and incubated overnight. Following a further wash, secondary antibody (goat anti-chicken 568 nm (Invitrogen, A11041, 1:1,000), streptavidin conjugate 647 nm (Invitrogen, 532357, 1:1,000), and goat anti-rabbit 488 nm (Invitrogen, A11034, 1:1,000)) was incubated for 1 hour. Following a final wash, slides were cured overnight with VECTASHIELD® Mounting Medium with DAPI (Vector Laboratories, H-1500) and imaged with a Leica DMI6000 B inverted fluorescent microscope.

To visualize EVs in astrocytes, primary murine astrocytes were plated at 2.5×10^5^ cells per well in 2 mL in a 6 well PLL coated plates. PKH67 dye (488 nm) stained the EVs according to the manufacturer’s protocol (Sigma, PKH67GL-1KT). EVs were added to astrocytes and incubated at 37°C with 5% CO_2_ for 2 hours. After incubation, supernatant was aspirated and cells washed once with 1X PBS, as described above. Primary antibody (rat GFAP (Invitrogen, 13-0300, 1:250)) and secondary antibody (donkey anti-rat 568 nm (Invitrogen, A78946, 1:1,000)) diluted in blocking buffer was added. Slides were cured overnight with VECTASHIELD® Mounting Medium with DAPI and imaged with a Keyence BZ-X all-in-one fluorescent microscope.

For blocking experiments endocytic pathway blockers were added 1 hour prior to the addition of PKH67 stained EVs. Astrocytes were incubated with both 25 μM EIPA (5-(N-Ethyl-Nisopropyl) amiloride) (Bio-techne, 3378/10) and 10 μM Cytochalasin D (Gibco, PHZ1063) to observe the EV uptake mechanism. Slides were imaged with a Leica DMI6000 B inverted fluorescent microscope.

### 2.8 Liquid Chromatography (LC)-Mass Spectrometry (MS)/MS

EVs were lysed in RIPA buffer (Thermo, 89901) containing protease inhibitors (Protease Inhibitor Mini Tablets, Thermo, A32955) at 400 μL of RIPA to 100 μL of EVs and incubated on ice for 15 minutes with occasional vortexing. EVs were passed through a 20-gauge needle 5 times and spun at 14,000 g for 15 minutes at 4°C. The supernatant was collected and transferred to an ultracentrifuge tube and washed with 2-3 mL of 8 M urea added and centrifuged at 4,400 rpm for 30 minutes at room temperature. This wash step was repeated three times. To each sample 20 μM dithiothreitol (DTT) (Bio-rad, 1610611) was added and incubated at 37°C for 1 hour. Following incubation 55 μM indoleacetic acid (IAA) (Sigma-Aldrich, I3750) was added and incubated for 30 minutes in the dark at room temperature. A buffer exchange to 50 mM ammonium bicarbonate (ABC) (Fisher Scientific, 02-002-270) was performed at 4,000 rpm for 20-40 minutes at room temperature. This step was repeated 4 times. To each sample 10 μg of Trypsin (Thermo Scientific, 20233) was added and incubated overnight for 16 hours at room temperature. Peptides were collected following centrifugation at 4,600 rpm for 40 minutes at room temperature and concentrated using a Speed-Vac. The peptides were desalted by a C18 ZipTip column (Thermo Scientific, 87782). The peptide solution was then dried by Speed-Vac and stored at −80°C until further LC-MS/MS analysis. Samples were harvested in triplicate.

For LC-MS/MS analysis, fractions were resuspended in 20 μL water with 0.1% formic acid (TCI America, 64-18-6) and separated by nano-LC followed by analysis by on-line electrospray tandem mass spectrometry. Experiments were performed on an EASY-nLC 1200 system (Thermo Fisher) connected to a quadrupole-Orbitrap mass spectrometer Orbitrap Fusion Tribrid Mass Spectrometry equipped with an EASY-Spray ion source (Thermo Fisher). From each peptide, 5 μL was loaded onto the trap column (Acclaim^TM^ PepMap^TM^ 100 C18 HPLC column, 75 μm x 25 cm, Thermo Scientific, 164946) with a flow of 10 μl/min for 3 min and subsequently separated on the analytical column (Acclaim^TM^ PepMap^TM^ C18) with a linear gradient, from 3% D to 40% D in 55 min. The column was re-equilibrated at initial conditions for 5 min. The flow rate was maintained at 300 nL/min and column temperature was maintained at 45°C. The electrospray voltage of 2.2 kV versus the inlet of the mass spectrometer was used. The Orbitrap Fusion Tribid Mass Spectrometry was operated in the data-dependent mode to switch automatically between MS and MS/MS acquisition. Survey full-scan MS spectra (m/z 375-1500) were acquired with a mass resolution of 60K, followed by fifteen sequential high energy collisional dissociation (HCD). The AGC target was set to 4.0E^5^, and the maximum injection time was 100 ms. MS/MS acquisition was performed in ion trap with the AGC target set to 3E^4^, and the isolation window at 1.6 m/z. Ions with charge states 2+, 3+, and 4+ were sequentially fragmented by HCD with a normalized collision energy (NCE) of 35%. Fixed first mass was set at 100. In all cases, one micro scan was recorded using a dynamic exclusion of 30 seconds.

LC-MS/MS raw data were processed and analyzed using MaxQuant (v 2.1.4.0) aligned to the TgondiiME49 (Toxo DB, version 68) and mouse (Uniport, ID UP000000589) genome databases. In particular, mass tolerances for precursor and fragment ions were 6 and 10 ppm, respectively, the minimum pep-tide length was 6 amino acids, and the maximum number of missed cleavages for trypsin was 2. LC-MS/MS validation of the peptide sequences for the most significant proteins were demonstrated (**Supplemental Figure 2**).

### 2.9 Western Blot

Primary astrocyte cultures were plated at 1×10^5^ cells/mL in 2 mL in 6-well plates. One day after plating Dexamethasone at 100 nM was added to stimulate GLT-1 expression. One day after Dexamethasone treatment experimental conditions were added: media alone, LPS/IFN*γ* (100 ng/mL and 100 U/mL), EVs from uninfected neurons (2 μg/mL), or EVs from infected neurons (2 μg/mL) and astrocytes incubated for 24 hours. EV concentration was based on publications and material (Antoniou et al., 2023; Solana-Balaguer et al., 2023). Cells were washed and lysed in RIPA buffer containing protease inhibitors, incubated on ice for 15 minutes with occasional shaking, followed by cell scraping and passage through a 20-gauge needle. The lysate was then spun at 14,000 g for 15 minutes at 4°C, supernatant collected and stored at −80°C. Protein concentration was measured by a BCA Protein Assay Kit (Thermo, 23250) according to the manufacturer’s protocol. Equal amounts of proteins (9 μg/mL) were separated by SDS-PAGE for 2 hours and transferred onto a 0.2 μm nitrocellulose membranes for 18.5 hours. Membranes were incubated in a blocking solution (0.1% PBST + 5% Milk) for 30 minutes at room temperature and then probed with primary antibodies (rabbit anti-β-actin (Abcam, AB8227, 1:1,000) and guinea-pig anti-GLT-1 (Millipore Sigma, AB1783, 1:500)) overnight at 4 °C. After washing three times for 5 min in 0.1% PBST (1X PBS + 0.1% Tween20), membranes were incubated with HRP-conjugated secondary antibodies (goat anti-rabbit HRP conjugate (BioRad, 170-5046, 1:5,000) and goat anti-guinea pig HRP conjugate (Invitrogen, A18769, 1:5,000)) for 1 h at room temperature. Membranes were washed five times for 10 minutes in 0.1% PBST. Signal was detected with the ECL Western blotting substrate (BioRad, 170-5061) and imaged on the Gel Doc XR+ System (BioRad). Blots were analyzed using ImageJ software. Additional MOIs (0.25, 0.50, 0.75, and 1.0) were also probed for β-actin and GLT-1 with equal amounts of protein added (7 μg/mL). Different concentrations of EVs (3 μg/mL, 0.50 μg/mL, and 0.10 μg/mL) were also probed for β-actin and GLT-1 with equal amounts of protein added (9 μg/mL).

### 2.10 RNA Analysis

EV RNA was isolated from neuronal supernatant after a series of ultracentrifugation methods, as mentioned previously. The EV RNA was then isolated through a membrane affinity exoRNeasy Midi Kit (Qiagen, 77144), according to the manufacturer’s protocol. RNA concentration and quality was determined by TapeStation RNA ScreenTape (Agilent Technologies) and High Sensitivity RNA Qubit (Thermo Fisher) at the UCR genomics core. Each condition was done in triplicate to obtain biological replicates.

Primary astrocytes were plated at 1×10^5^ cells per well in 2 mL in 6 well plates with the same conditions listed under the Western Blot section. After 24 hours of incubation the wells were washed once with 1X PBS and lysed with 350 µl of Buffer RLT (Qiagen RNeasy Mini Kit, 74104). Cells were harvested and passed through a 20-gauge needle 5 times. Following lysis, the RNeasy Mini Kit was utilized following the manufacturer’s protocol. RNA concentration and quality was determined by TapeStation RNA ScreenTape (Agilent Technologies) at the UCR genomics core. Samples with a RIN (RNA Integrity Number) > 9 were chosen. Each condition was conducted in triplicate to obtain biological replicates.

For EV RNA sequencing library construction, quality control, and small RNA sequencing (RNA-Seq) were performed by the UCR Genomics Core. Libraries were constructed with the Qiagen QIAseq miRNA Library Kit (331502) along with the NEBNext Library Quant kit for Illumina (E7630S) for additional quality control. Samples were sequenced on the Illumina NextSeq2000 platform with SE100 around 100 million reads per sample. Raw data in fastq format was processed through CLC genomics workbench (v24.0.2) to remove UMIs (unique molecular identifiers) and aligned to miRbase (v22) mouse genome. Simultaneously, QC and GC content of the clean reads was calculated. All downstream analyses were based on high quality clean data. Differential expression analysis of biological replicates was performed using the DESeq2 R package (v1.20.0). Feature counts were normalized using the normalization function in the DESeq2 package. miRNAs with a p-value <=0.05 and a fold change (FC) >0, found by DESeq2, were assigned as differentially expressed.

For astrocyte RNA sequencing library construction, quality control, and bulk RNA-Seq were performed by UCSD Genomics Core with 25 ng RNA used for each sample. Libraries were generated using the Illumina Stranded mRNA Prep according to the manufacturer’s protocol. Constructed libraries were sequenced on the Illumina NovaSeqXPlus LH00444 (240223_LH00444_0061_B223HCLLT4) platform with PE150 around 25 million reads per sample. Raw data in fastq format was processed through FastQC software (v0.11.9) and clean reads were obtained by the removal of reads containing adapters, poly-N and low-quality through Trim Galore software (v0.6.7). Simultaneously, QC and GC content of the clean reads was calculated. All downstream analyses were based on high quality clean data. Reads were aligned to the mouse genome (Gencode, GRCm39, vM34) using STAR (v2.7.11a) and featureCounts (v1.5.0-p3) was used to count the number of reads mapped to each gene. Differential expression analysis of biological replicates was performed using the DESeq2 R package (v1.20.0). Feature counts were normalized using the normalization function in the DESeq2 package. Genes with a p-value <=0.05 and a fold change (FC) >0, found by DESeq2, were assigned as differentially expressed. Gene Ontology (GO) enrichment analysis of differentially expressed genes was performed by the clusterProfiler R package, in which gene length bias was corrected. GO terms with p-values less than 0.05 were considered significantly enriched by differential expressed genes.

### 2.11 Data Analysis

Data was plotted using GraphPad Prism 9.0 software, replicates were analyzed using mean and standard deviation, and statistical significance was determined using an unpaired t-test or one way-ANOVA (p<0.05).

## 3 Results

### 3.1 *T. gondii* infection of neurons decreases the production of EVs

To determine if infection of neurons alters EV production, EVs were isolated from primary cortical neurons infected with *T. gondii.* Toxoplasma spontaneously forms cysts in neurons (Halonen et al., 1996; Mouveaux et al., 2021), however we have found that too many parasites overwhelm the cultures. To determine the maximum number of infected neurons with the least amount of neuronal damage, four different multiplicity of infections (MOIs) (0.25, 0.50, 0.75, and 1.0) were tested. Primary cortical neurons were infected with their respected MOIs, 8 days after culture to ensure full development and confluency of neurons pre-infection, and images were taken 3 days post infection (d.p.i). Neuronal morphology was confirmed with MAP2 (microtubule associated protein 2) staining of dendrites, and NeuN (neuronal nuclear antigen) staining of the nucleus. As previously published, upon infection of neurons, Toxoplasma spontaneously formed cysts (Mouveaux et al., 2021; Wohlfert et al., 2017) (**Figure 1A**). Cyst generation was confirmed with DBA (*Dolichos biflorus* agglutinin) that binds to sugars of the cyst wall (**Figure 1A**; **Supplemental Figure 1A**). A decrease in dendrite number was observed as MOI increased (**Supplemental Figure 1B-C**). To maximize the number of infected neurons in culture while optimizing neuronal health, an MOI of 0.50 was chosen for the remaining experiments. Infection with a 0.50 MOI did not alter the morphology or percentage of NeuN and MAP2 positive neurons, indicating no neuronal toxicity (**Figure 1**). *In vitro* studies of *T. gondii* have shown cysts to be present in both the dendrite and soma of neurons, which is consistent with our data (Halonen, 2023; Mendez et al., 2018, 2021; Mendez & Koshy, 2017) (**Figure 1C**).

**Figure 1:**
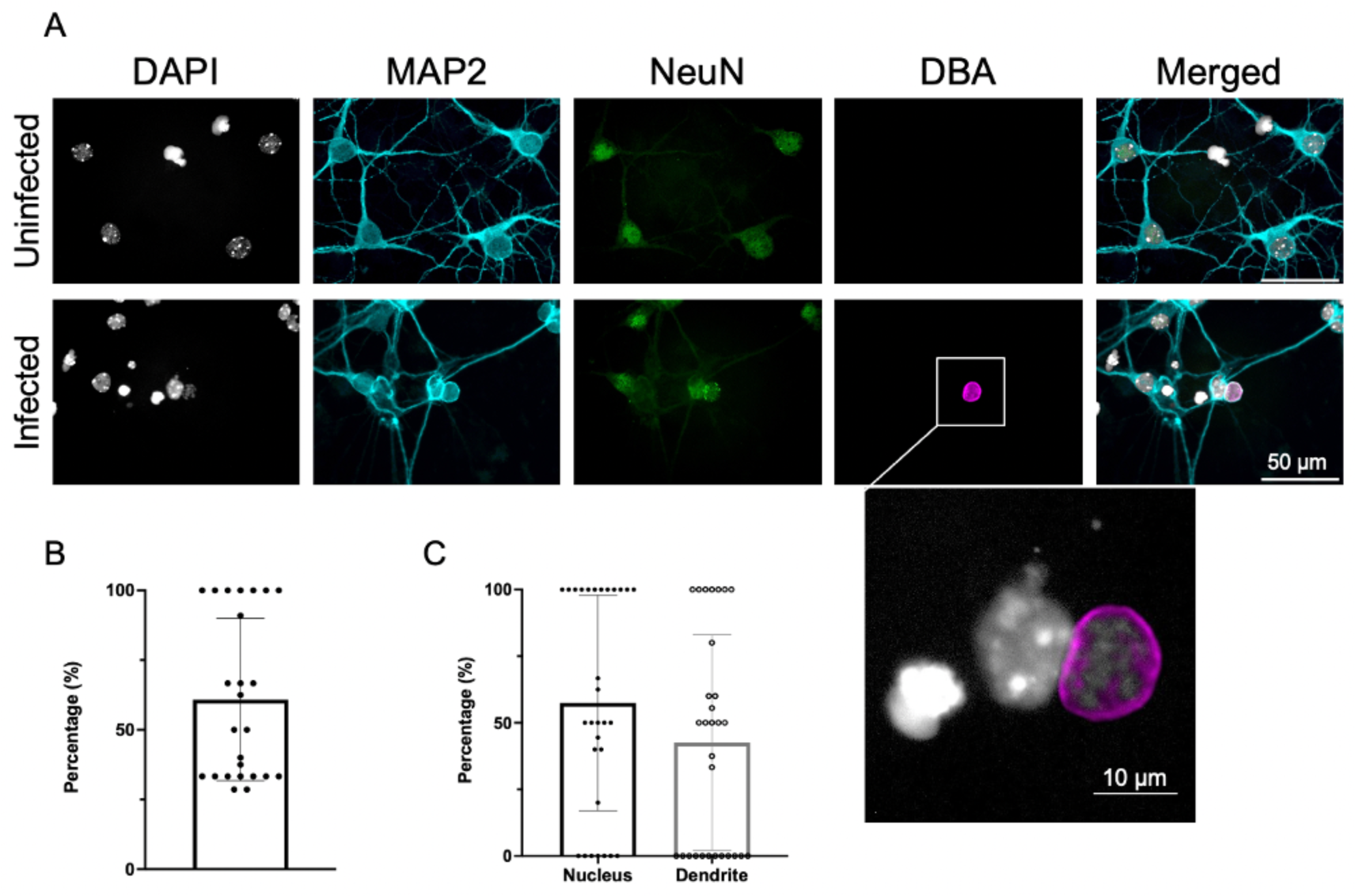
Primary cortical neurons infected with Toxoplasma. Infection with a MOI (multiplicity of infection) of 0.5. **A)** Images taken three days post infection. DAPI stains the nucleus, MAP2 stains the perikarya and dendrites on neurons, NeuN stains the nucleus of the neurons, DBA stains the sugars of the cyst wall. In the infected neurons there is overlap of the cyst and neuron. Scale bar at 50 µm. **B)** Quantification of the percentage of infected neurons. The number of cysts and neuronal nuclei were counted and the percentage of infection was calculated. Around half of the neurons were infected with *T. gondii,* which is consistent with the MOI of 0.5. **C)** Quantification of the percentage of cysts located in the nucleus of the neuron versus the dendrite of the neuron (Unpaired t-test, n (Nucleus) = 30, n (Dendrite) = 30, Nucleus vs. Dendrite p value = 0.1586).

EVs were harvested from neurons at day 3 post infection. To confirm EV characteristics, transmission electron microscopy (TEM) was conducted to determine size or morphological changes upon infection. TEM confirmed the circular morphology of EVs from both uninfected and infected neurons, consistent with previous publications (Barranco et al., 2019; Kotrbová et al., 2019; Rikkert et al., 2019; Veraguas et al., 2021) (**Figure 2A**). Quantification of EV size demonstrated no significant difference between EVs from uninfected and infected neurons, and both were within EV range, with uninfected averaging at 76 nm in diameter and infected averaging at 77 nm in diameter (Van Der Pol et al., 2014; Vestad et al., 2017) (**Figure 2B**).

**Figure 2:**
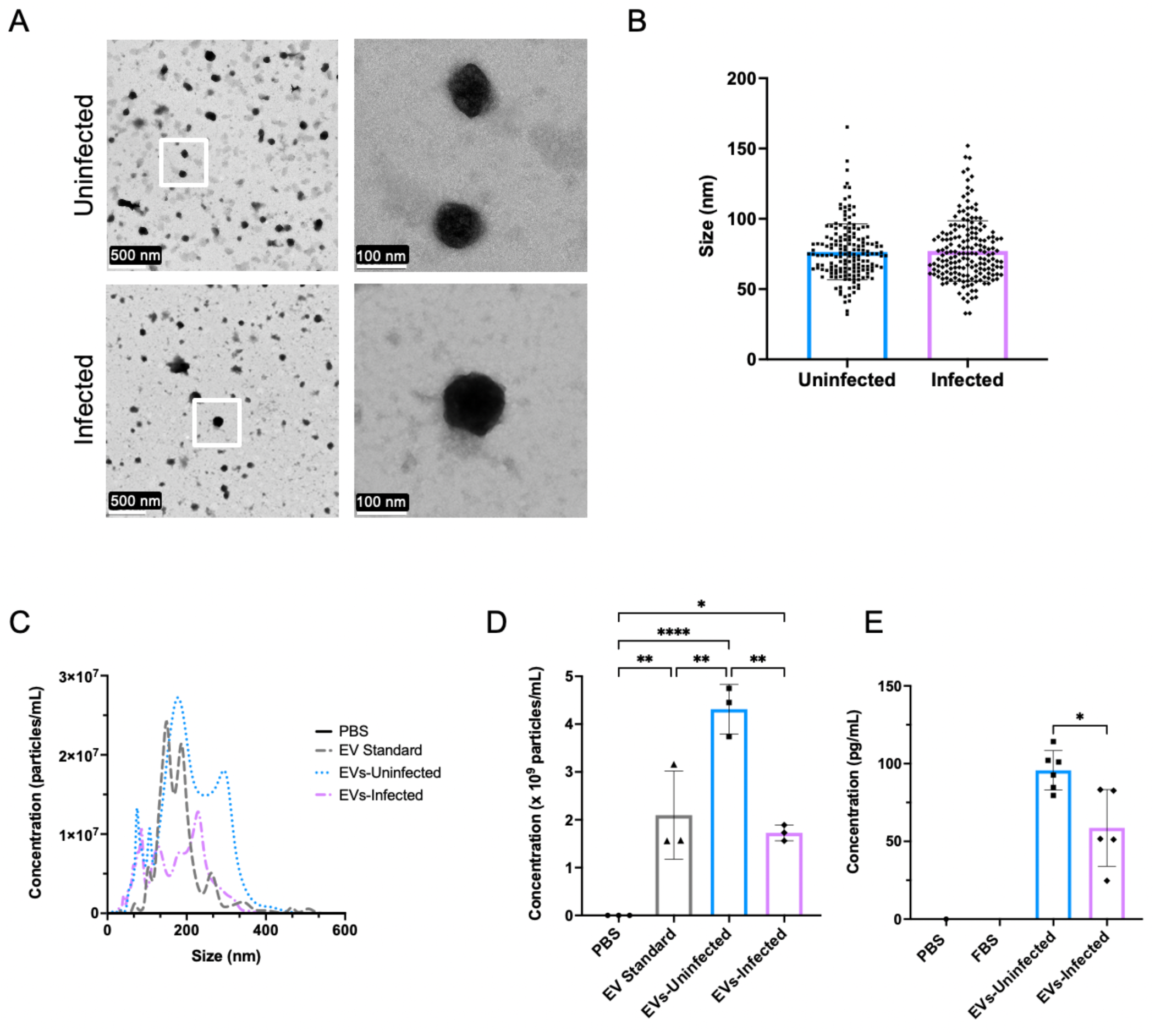
Toxoplasma infection leads to a decrease in the concentration of EVs. **A)** Transmission Electron Microscopy (TEM) of EVs isolated from uninfected and infected neurons. Right hand panels are zoomed in images of areas outlined by the white box in the corresponding left panels. **B)** Quantification of the size of EVs from TEM images (Unpaired t-test, n (Uninfected) = 169, n (Infected) = 179, Uninfected vs. Infected p value = 0.7691). **C)** Nanoparticle Tracking Analysis (NTA) of particles from PBS, EV Standard, EVs isolated from uninfected neurons, and EVs isolated from infected neurons. **D)** Concentration of nanoparticles in neuronal supernatants as determined by NTA and CD63 ELISA after centrifugation and ExoEasy Kit (One-way ANOVA, n = 3 EV Standard vs. EVs-Uninfected p value = 0.0042, EV Standard vs. EVs-Infected p value = 0.8309, PBS vs. EVs-Uninfected p value = <0.0001, PBS vs. EVs-Infected p value = 0.0177, PBS vs. EV Standard p value = 0.0059, EVs-Uninfected vs. EVs-Infected p value = 0.0016). **E)** CD63 ELISA (Unpaired t-test, n (PBS) = 1, n (FBS) = 0, n (EVs-Uninfected) = 6, n (EVs-Infected) = 5, EVs-Uninfected vs. EVs-Infected p value = 0.0104).

Nanoparticle tracking analysis (NTA) was performed to determine the average size and concentration of EVs. Phosphate buffered saline (PBS) was used as a negative control (Kusuma et al., 2018; Witwer et al., 2013). EVs from C57BL/6 mouse embryo fibroblasts were used as a positive control at a known particle concentration (STEMON, 1×10^9^ particles/mL). NTA allowed for visualization of the distribution of particles in the control and experimental groups (**Figure 2C**). NTA quantification of EVs from uninfected and infected neurons demonstrated a significant decrease in particle concentration following infection (p<0.005) (**Figure 2D**). Since NTA measures all particles and is not specific to EVs, a CD63 ELISA was conducted (Bachurski et al., 2019). CD63 is one of the three tetraspanin proteins, along with CD81 and CD9, commonly found on the surface of EVs and is widely used as an identifying EV marker (Ter-Ovanesyan et al., 2023; Théry et al., 2018). CD63 quantification confirmed the presence of EVs from neuronal samples and that infection with *T. gondii* decreases CD63+ EV concentration (p<0.05) (**Figure 2D**). Overall, these experiments determined that following Toxoplasma cyst infection of murine cortical neurons there is a significant decrease in neuronal production of EVs.

### 3.2 *T. gondii* infection alters the protein content of EVs

EVs contain multiple proteins, many of which are involved in the adhesion of the vesicle to the target cell (Théry et al., 2002) but many that can also reveal function and act as biomarkers of disease (Karnati et al., 2019). To determine if Toxoplasma infection alters the content of EVs, analysis of proteins via liquid chromatography (LC)-mass spectrometry (MS)/MS was conducted on uninfected and infected neuronal derived EVs. A principal component analysis (PCA) plot was generated to determine the grouping and relationship between EV samples from uninfected (blue) and infected (purple) neurons, demonstrating that protein signatures were different between the two samples (**Figure 3A**). When comparing protein similarities between the two groups there are 183 proteins that were consistently found in EVs from both uninfected and infected neurons. Only 17 proteins were uniquely identified in EVs from uninfected neurons, however, 101 proteins were found specifically in EVs from infected neurons (**Figure 3B**). Of all proteins only 20 were significantly differentially regulated and consisted of 6 proteins found only in EVs from infected neurons, and the remaining 14 found in both groups. Proteins significantly altered following infection were aligned to the mouse genome and a volcano plot was generated to visualize infection dependent protein changes (**Figure 3C**). There were 12 proteins significantly upregulated (red) in EVs from infected neurons, and 8 significantly downregulated (blue) (**Figure 3C**; **Supplemental Table 1**). Peptide verification was performed for three different protein sequences (**Supplemental Figure 2**). Highly upregulated proteins included vimentin (Vim), tubulin beta-4B chain (Tubb4b) and endoplasmin (Hsp90b1). These proteins have associations with the interferon response, innate immunity, stress response, and structural support (**Supplemental Table 1**). Tubb4b and Hsdp901 are commonly found in EVs, with Hsp90b1 being an identifying marker for isolated EVs (Dozio & Sanchez, 2017; Kurg et al., 2022). Vim, a component of the cytoskeleton and vital for the regulation of intracellular signaling pathways (Liu et al., 2022) was expressed almost 8-fold higher in EVs from infected neurons. In contrast to the upregulation of immune associated pathways, the top 3 significantly downregulated proteins were involved with neuronal migration and growth (e.g. Reln, Chl1, Aplp1) (**Supplemental Table 1**). Reelin (Reln) is the most downregulated protein in EVs following infection and is linked to neuromodulation by protecting the brain against neurodegeneration and controlling synaptic plasticity (Joly-Amado et al., 2023). Neuronal cell adhesion molecule L1-like protein (Chl1) and amyloid beta precursor like protein 1 (Aplp1) are involved in dendritic spine pruning and synaptic transmission (Mohan et al., 2019; Vnencak et al., 2015). Gene ontology (GO) was performed on the significant proteins to help identify important functional properties related to protein changes. Thus, upon infection EV protein content suggests a downregulation of synapse organization and migration, and an upregulation in cell adhesion, binding, and intracellular signaling (**Figure 3D**).

**Figure 3:**
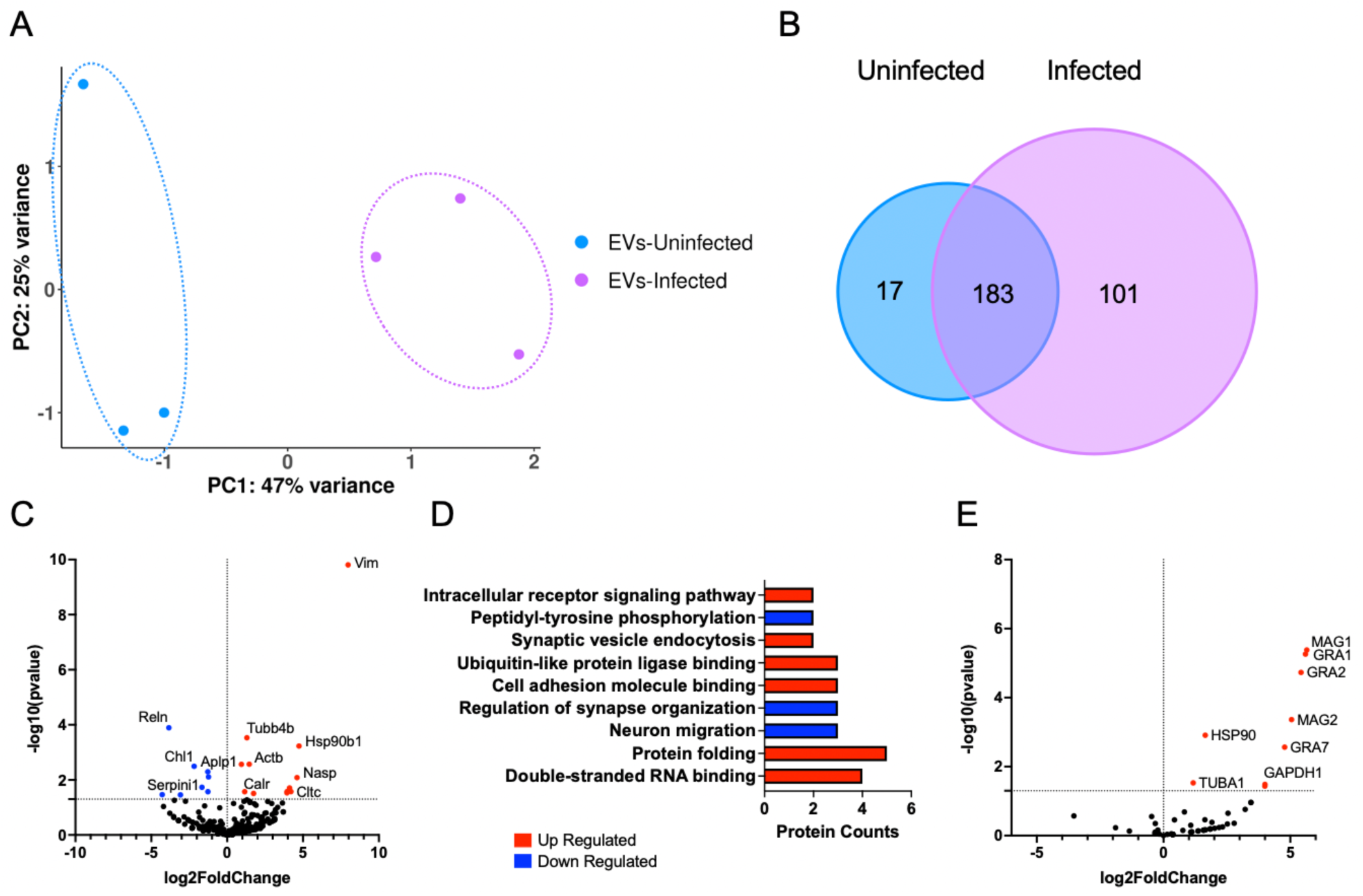
Toxoplasma infection leads to changes in EV protein content. **A)** PCA (principal component analysis) of EVs from uninfected neurons (blue) and infected neurons (purple) after LC-MS/MS protein analysis. **B)** Venn Diagram depicting the protein alterations following infection. **C)** Volcano plot of protein changes aligned to the mouse genome. Above −log10(0.05) was considered significant. From the volcano plot 12 proteins are upregulated (red), 8 proteins are downregulated (blue), and 281 proteins are not changed (black). **D)** GO analysis of Uninfected vs. Infected significant EV proteins. Red boxes indicate upregulated GO terms and blue boxes represent downregulated GO Terms. P-value increase going up the y-axis. **E)** Volcano plot comparing significant protein alterations following infection when comparing to the Toxoplasma genome. Anything above −log10(0.05) was considered significant. From the volcano plot 9 proteins are upregulated (red), 0 proteins are downregulated (blue), and 48 proteins are not changed (black).

Toxoplasma possess sophisticated machinery to inject parasite proteins into host cells during and in the absence of host cell invasion, altering cell signaling events (Hunter & Sibley, 2012; Koshy et al., 2012). In addition, the parasite, itself a eukaryotic cell, is capable of EV production (Długońska & Gatkowska, 2016; Maia et al., 2021; Quiarim et al., 2021). Thus, collection of EVs from infected neurons has the potential to contain parasite EVs and/or proteins excreted by the parasite and subsequently incorporated into host EVs. To identify any parasite specific proteins, alignment of protein sequences was also made to the Me49 strain of *T. gondii* (**Figure 3E**). This demonstrates the presence of 9 Toxoplasma proteins contained in neuronal EVs following infection. Some of these proteins are housekeeping proteins often incorporated into EVs (e.g, HSP90, TUBA1, GAPDH1) (Karnati et al., 2019), but others are *T. gondii* specific proteins known to be secreted effectors (e.g. MAG1, GRA1, GRA2, MAG2, and GRA7) involved in cyst wall formation and host cell manipulation (Ihara & Nishikawa, 2021; Nam, 2009; Tu et al., 2020) (**Figure 3E**). In addition, two cyst-wall specific proteins, MAG1 and MAG2, were present in EVs from infected neurons (Cruz Camacho et al., 2023; Nam, 2009) (**Supplemental Table 2**). These results demonstrate that infection of neurons with Toxoplasma alters EV content including the addition of parasite proteins.

### 3.3 Infection alters the miRNA content of EVs

The cargo of EVs can be diverse and includes DNA, RNA, and protein. Micro RNAs (miRNAs) can alter gene expression by post-transcriptional silencing or via chromatin modifications to silence or activate genes (O’Brien et al., 2018). Previous work has demonstrated miRNAs (miR), specifically miR-124, from neuronal EVs regulate GLT-1 expression on astrocytes (Morel et al., 2013; Pinto et al., 2017). To determine if infection of neurons also alters the miRNA content of EVs, miRNA-Sequencing was conducted of EVs harvested from uninfected and infected neurons. miRNA transcript analysis determined distinct separation of EVs from uninfected (blue) and infected (purple) neurons, although with a broader variation of EVs from uninfected neurons. This indicates a parasite driven alteration in miRNA content (**Figure 4A**). Significantly expressed miRNA z-scores were normalized in order to obtain a heatmap to visualize the enriched miRNAs and observe similarities between the triplicates of each group and differences following infection (**Figure 4B**). When aligned to the mouse genome, 264 miRNAs were significantly differentially expressed with 8 miRNAs upregulated (red) and 6 miRNAs downregulated (blue) in EVs from infected neurons (**Figure 4C**). While some of the identified miRNAs do not have predicted gene targets and known functions, there are many that have been characterized. Of the downregulated miRNAs, both miR-3473b and miR-1224-5p have been reported to enhance inflammation and chemokine release in the brain (Feng et al., 2021; X. Wang et al., 2018) (**Supplemental Table 3**). In addition, miR-324-5p regulates excitatory synapse structure and function (Parkins et al., 2023, 2023) and miR-466i-5p is predicted to induce neuronal apoptosis (Zhu et al., 2022). Of the upregulated miRNAs, miR-196a-5p, miR-199a-3p, miR-143-3p, miR-21a-5p have all been linked to an anti-inflammatory response through the inhibition of the NF-κB pathway (Duan et al., 2022; Huang et al., 2014; Peng et al., 2020; Saadh et al., 2023; Y. Wang et al., 2020; Xiong et al., 2019; Ye et al., 2018) (**Supplemental Table 3**), with the exception of miR-29a-3p which exhibits a pro-inflammatory response through the phosphorylation of Akt and downstream NF-κB activation (Tang et al., 2017). Of note, no significant difference was observed in miR-124 (miR-124-5p), responsible for GLT-1 regulation (Men et al., 2019; Morel et al., 2013; Ponomarev et al., 2011), suggesting other mechanisms are responsible for the reduction in astrocytic GLT-1. Thus, infection of neurons with Toxoplasma alters the small RNA content of EVs, specifically a downregulation of pro-inflammatory miRNAs, and an upregulation of anti-inflammatory miRNAs, however, does not support an infection-induced change in GLT-1 through miRNA-124 regulation.

**Figure 4:**
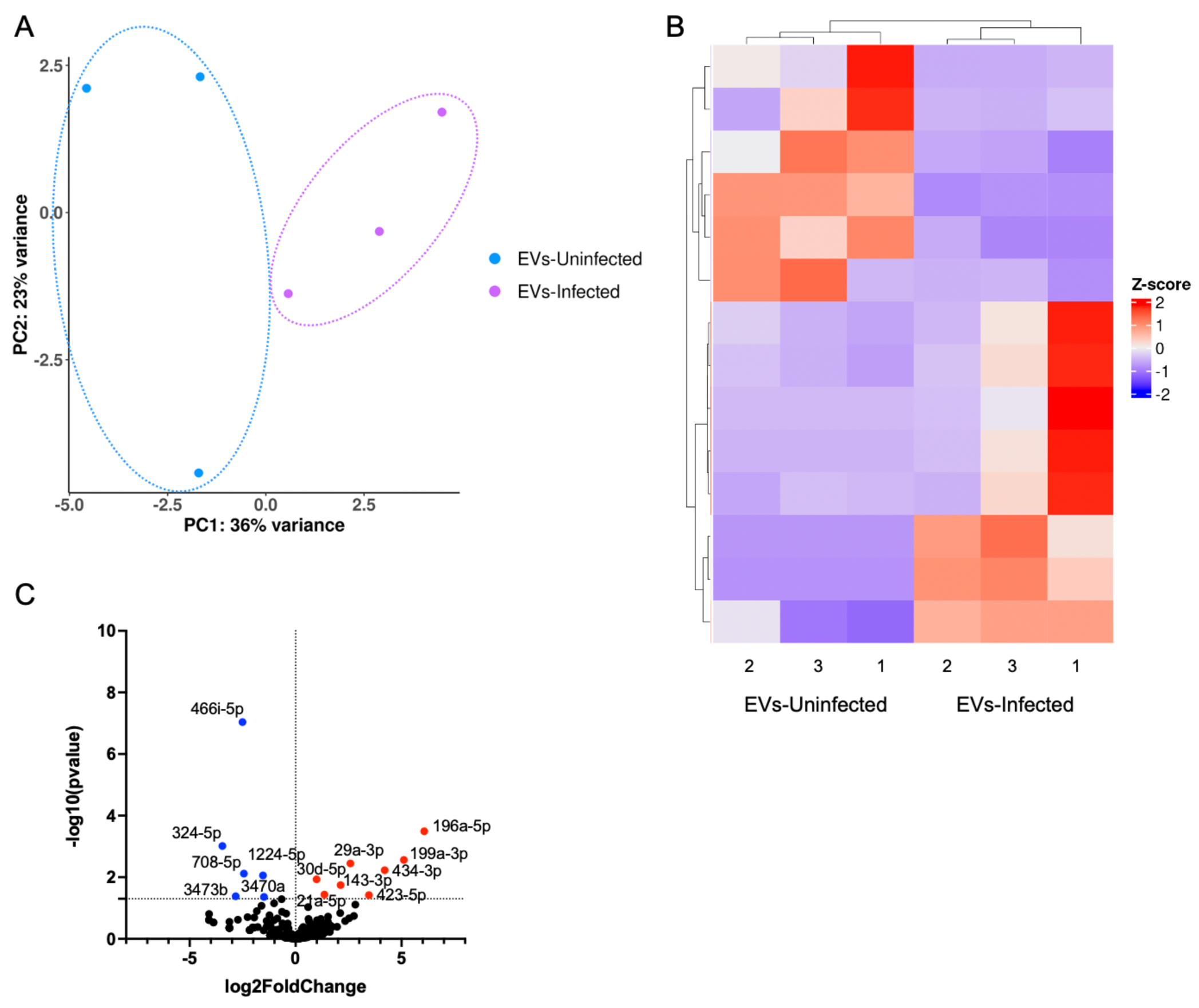
Infection alters neuronal EV miRNA content. **A)** PCA of EVs from uninfected neurons (blue) and infected neurons (purple) after miRNA Sequencing. Three replicates per group. **B)** Heat map of significant differentially enriched miRNAs after the addition of EVs from Uninfected EV and Infected EVs. Each sample was conducted in triplicate. Each count was normalized, and the Z-score was mapped with columns and rows being clustered. **C)** Volcano plot of differentially enriched miRNA comparing neuronal EV after infection. Anything above −log10(0.05) was considered significant. From the volcano plot 8 miRNA are upregulated (red), 6 miRNA are downregulated (blue), and 250 miRNA are not changed (black).

### 3.4 EVs from infected neurons are taken up by astrocytes

During Toxoplasma infection of an immunocompetent host, the dominant infected cell is the neuron. However, infection has the potential to have a much broader impact on the neurological landscape via injection of parasite proteins into cells that it does not invade (Mendez et al., 2021) and as a result of the significant immune response necessary to control infection. Neurons have an intimate relationship with astrocytes and astrocytes can absorb neuronal-derived EVs, resulting in an alteration in their gene expression, protein levels, and immune function (Men et al., 2019; Morel et al., 2013; Ogaki et al., 2021; Ponomarev et al., 2011). To determine if infection-induced EVs alter astrocyte gene expression, EVs from uninfected and infected neurons were incubated with primary murine astrocytes. Isolated EVs were stained with PKH67, a green lipophilic dye, to visualize uptake and location of EVs (Y. Li et al., 2018; Yin et al., 2020). PKH67 labeled EVs colocalized in GFAP+ astrocyte cultures independent of the infection conditions of the neurons from which they were derived (**Figure 5**). EVs could be seen in the cytoplasm, often associated with vacuolar structures (**Figure 5A/B**). Interestingly, EVs could also be seen in the nucleus of astrocytes (**Figure 5A/B**). To further confirm the intracellular localization of EVs, confocal images were reconstructed in a 3D model to determine the z-planes in which EVs are found. When decreasing in z-plane the EVs come into and out of focus within the cell (**Figure 5B**; **Supplemental Figure 3**). Our data suggests a significant alteration in protein content of EVs from infected neurons which may influence EV uptake by target cells. Quantification of EV localization was conducted to determine if the infection status of the neurons altered EV uptake and cellular localization within astrocytes. Infection did not lead to a difference in the proportion of EVs taken up by astrocytes (**Figure 5C**), nor did it alter their nuclear or cytoplasmic localization (**Figure 5D-E**).

**Figure 5:**
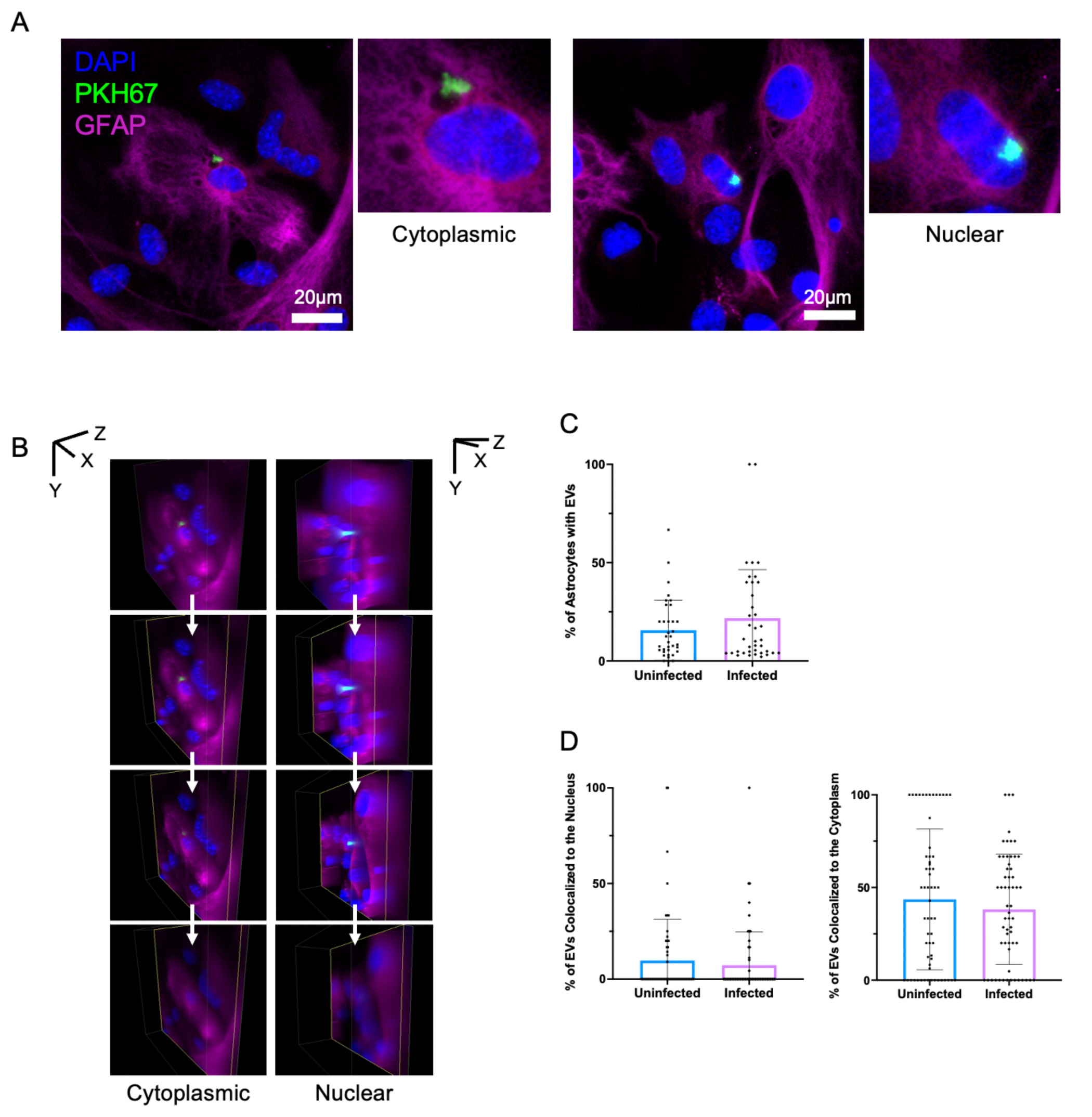
Astrocytes take up neuronal EVs. **A)** Fluorescent microscopy of astrocytes after the addition of EVs from uninfected and *T. gondii* infected neurons indicating the location of EVs in the cytoplasm versus nucleus. Scale bar: 20µm. **B)** 3D reconstruction of z-stacks to identify the location of the EVs. A decrease in a pitch of 0.4µm for uninfected and 0.2µm for infected, to depict the uptake of EVs by astrocytes. **C)** Quantification of the percentage of astrocytes that contain EVs (Unpaired t-test, n (Uninfected) = 35, n (Infected) = 38, Uninfected vs. Infected p = 0.2081). **D)** Quantification of the percentage of EVs that are colocalized with the nucleus of the astrocytes (Unpaired t-test, n (Uninfected) = 61, n (Infected) = 59, Uninfected vs. Infected p = 0.4900). Quantification of the percentage of EVs that are colocalized with the cytoplasm of the astrocytes (Unpaired t-test, n (Uninfected) = 61, n (Infected) = 59, Uninfected vs. Infected p = 0.3853).

Having demonstrated astrocytes can uptake EVs from both uninfected and infected neurons we were interested in the EV uptake mechanism that is poorly understood in astrocytes. Previous studies have indicated that microglia take up oligodendrocyte derived EVs through the macropinocytosis pathway (Fitzner et al., 2011; Pantazopoulou et al., 2023). To better understand the mechanism of EV uptake by astrocytes, endocytic pathways were blocked and EV uptake measured. Two different endocytic pathway inhibitors were added to astrocyte cultures, 1 hour prior to the addition of EVs (**Supplemental Figure 4**). When blocking macropinocytosis by both EIPA and cytochalasin D, EV uptake was still observed in astrocytes from both EV groups. This data suggests that neuronal derived EVs are not taken up by astrocytes through macropinocytosis and alternative mechanisms such as clathrin mediated or caveolin endocytosis need to be considered (**Supplemental Figure 4B**). Altogether, these data demonstrate that EVs from uninfected and infected neurons are internalized by astrocytes via a micropinocytosis-independent pathway and localize to both the cytoplasm and astrocytic nucleus.

### 3.5 EVs alter astrocyte gene expression and GLT-1 protein expression in an infection-dependent manner

Our previous study demonstrated a reduction in astrocytic GLT-1 in the brain of infected mice and that this reduction was, at least in part, responsible for neuronal pathology. GLT-1 loss was not directly correlated to areas of infected astrocytes (David et al., 2016). Previous work supports the concept that neurons communicate and can regulate astrocyte function via EV release, including the expression of GLT-1 (Men et al., 2019; Morel et al., 2013; Ponomarev et al., 2011). To test this and determine if infection of neurons can alter this avenue of communication, EVs from uninfected and infected neurons (2 μg/mL) were incubated with astrocytes and astrocyte gene expression measured via RNASeq. Astrocytes with the addition of media alone were used as a baseline control. *In vitro* derived cultured astrocytes do not express GLT-1 therefore Dexamethasone (Dex) was added to increase GLT-1 expression, acted as a positive control for GLT-1 and to determine the ability of EVs from infected neurons to downregulate GLT-1 (Carter et al., 2012; Zschocke et al., 2005). In addition, astrocytes stimulated with LPS and IFN*γ*, known to induce a significant pro-inflammatory response in the astrocyte, acted as a positive control for known changes in gene expression (Hamby et al., 2012). Analysis of overall gene expression in replicates and stimulations via PCA verified groupings and our negative and positive controls (**Figure 6A**). When focusing only on the effects of the addition of EVs from uninfected and infected neurons, PCA confirmed replicates clustered together and were distinct from each other (**Figure 6B**). This indicates the addition of EVs produced by infected neurons significantly alters astrocyte gene expression compared to EVs from uninfected neurons.

**Figure 6:**
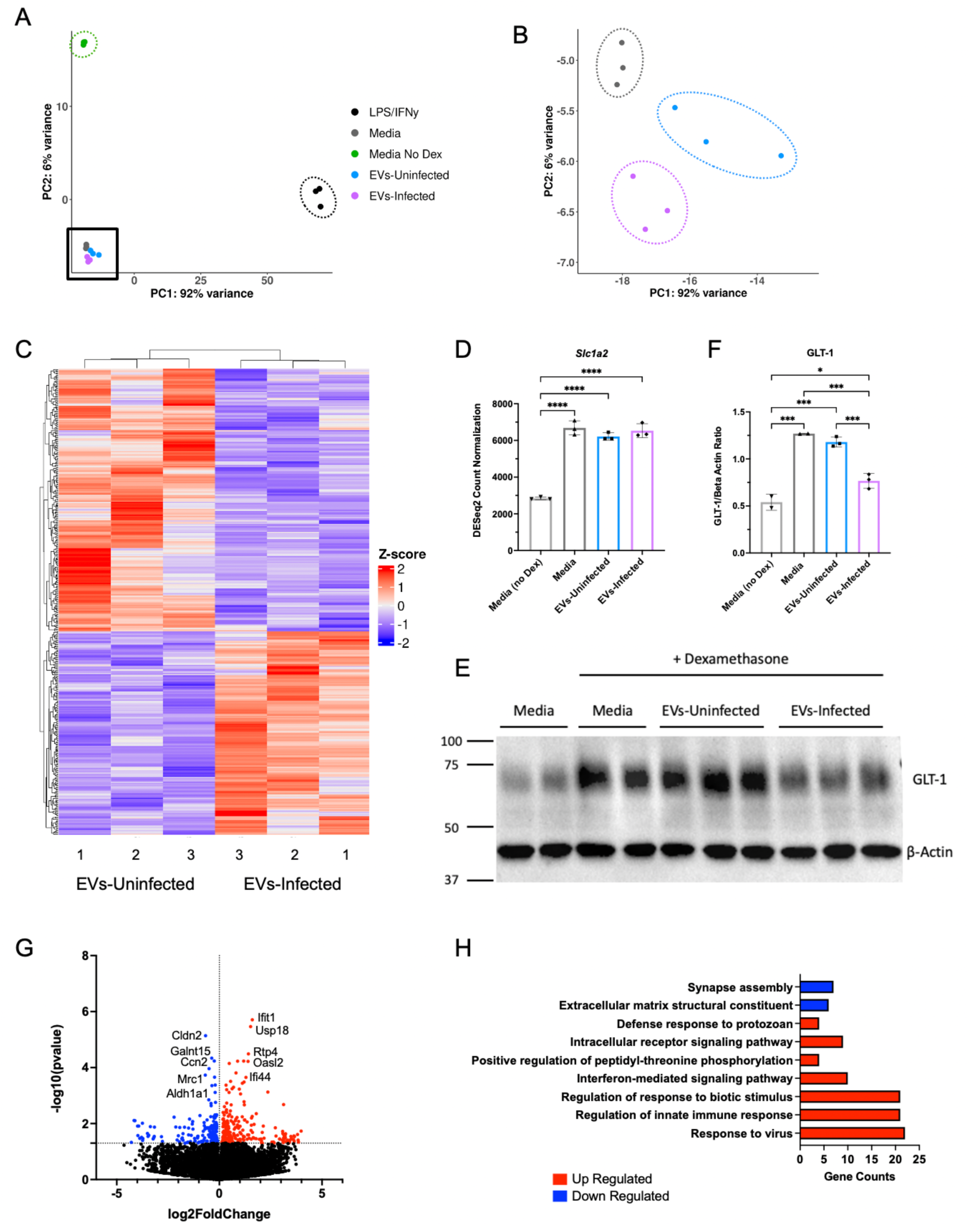
EVs from infected neurons alter astrocyte gene expression and decrease GLT-1 protein expression. **A)** PCA of astrocytes after 24 hours with the addition of LPS/IFNγ (black), cell culture media (gray), cell culture media without dexamethasone (green), EVs from uninfected neurons (blue) and EVs from infected neurons (purple). **B)** Zoomed in PCA plot focusing on Uninfected EVs, and Infected EVs **C)** Heat map of significant astrocyte DEGs after the addition of EVs from Uninfected EV and Infected EVs. Each sample was conducted in triplicate. Each count was normalized, and the Z-score was mapped with columns and rows being clustered. **D)** Slc1a2 gene expression levels of astrocytes after the addition of astrocyte media without dexamethasone, astrocyte media, EVs from uninfected neurons, and EVs from infected neurons. Comparison of Slc1a2 expression (gene for GLT-1), using DESeq2 Normalized counts. (One-way ANOVA, n= 3, Media (no Dex) vs. Media p value = <0.0001, Media (no Dex) vs. EVs-Uninfected p value = <0.0001, Media (no Dex) vs. EVs-Infected p value = <0.0001, Media vs. EVs-Uninfected p value = 0.2734, Media vs. EVs-Infected p value = 0.9142, EVs-Uninfected vs. EVs-Infected p value = 0.5726). **E)** Western blot for GLT-1 (62kD) protein expression. β-Actin (42 kDa) used as loading control. **F)** Quantification of Western Blot bands using Fiji (ImageJ). A ratio of the pixels (GLT-1/***β***-Actin) was taken and plotted. (One-way ANOVA, n (Media without Dex.) = 2, n (Media) = 2, n (EVs-Uninfected) = 3, n (EVs-Infected) = 3, Media without Dex. vs. Media p value = 0.0001, Media without Dex. vs. EVs-Uninfected p value = 0.0002, Media without Dex. vs. EVs-Infected = 0.0325, Media vs. EVs-Uninfected p value = 0.4825, Media vs. EVs-Infected p value = 0.0006, EVs-Uninfected vs. EVs-Infected p value = 0.0010). **G)** Volcano plot of DEGs comparing astrocytes RNA after the addition of EVs from uninfected neurons vs. the addition of EVs from infected neurons. Anything above −log10(0.05) was considered significant. From the volcano plot 227 genes are upregulated (red), 176 genes are downregulated (blue), and 31,135 genes are not changed (black). **H)** GO analysis of Uninfected EV vs. Infected EV DEGs. Red boxes indicate upregulated GO terms and blue boxes represent downregulated GO Terms. P-value increase going up the y-axis.

A comparison of DESeq2 normalized counts between the four groups confirmed the upregulation of GLT-1 gene expression (*Slc1a2*), with the addition of Dex (p<0.0001) and EVs (**Figure 6D**). However, there is no difference in GLT-1 gene expression between the addition of uninfected versus infected EVs (**Figure 6D**). In addition to inhibition of transcription, GLT-1 can also be regulated by post-translational modifications (Peterson & Binder, 2019). To determine if EVs from infected neurons alter GLT-1 protein expression a western blot for GLT-1 was conducted using the same conditions as before (**Figure 6E**). Quantification of the normalized bands (GLT-1/β-Actin) identified a significant decrease in GLT-1 protein intensity after the addition of EVs from infected neurons (p<0.005) (**Figure 6F**). Further experiments indicated the decrease in GLT-1 protein expression was not dependent on the concentration of EVs added or the MOI of the parasite, leading us to hypothesize that there may be post-translational modifications at play and the decrease is dependent on EV content (**Supplemental Figure 5-6**).

To expand our understanding of the effect of neuronal infection on astrocyte gene transcription, significantly expressed DEG z-scores were normalized and visualized on a heatmap to determine genes that are highly enriched in astrocytes after the addition of EVs from uninfected versus infected neurons. This revealed a similarity within biological replicates and striking patterns of genes that were significantly different (**Figure 6G**). Significance was determined by having a p-value greater than the negative log of 0.05 and having a log2FC (fold change) greater or less than 0. Analysis of differentially expressed genes (DEGs) with a p-value<0.05 were plotted to identify which genes were upregulated and downregulated in response to infection (**Figure 6G**). While a majority of the genes were not altered (31,135 genes), there were 227 genes upregulated and 176 genes downregulated following addition of EVs from infected neurons. The most significantly upregulated genes (e.g. *Ifit1*, *Usp18*, *Rtp4*) are all involved in a pro-inflammatory innate immune response. Interferon-induced protein with tetratricopeptide repeats 1 (*Ifit1*) induces type I interferon signaling (Reynaud et al., 2015), while the ubiquitin specific peptidase 18 (*Usp18*) and receptor transporter protein 4 (*Rtp4*) negatively regulates type I interferon signaling (He et al., 2020; Hou et al., 2021; Sui et al., 2024; Yim et al., 2016). The top significantly downregulated genes are associated with cellular communication (e.g. *Cldn2*, *Galnt15*, *Ccn2*) (**Supplemental Table 3**). These genes act in cellular binding and adhesion and all are important for allowing the attachment and entry of molecules, with claudin 2 (*Cldn2*) allowing for paracellular permeability (Weber & Turner, 2017); polypeptide N-acetylgalactosaminyltransferase 15 (*Galnt15*) predicted to enable carbohydrate binding (Valdés-Hernández et al., 2024); and cellular communication network factor 2 (*Ccn2*) permits cell adhesion (Hall-Glenn & Lyons, 2011), indicating that the addition of EVs from infected neurons is reducing the adhesive properties of astrocytes. GO analysis confirmed the upregulation of immune responsiveness of astrocytes including the upregulation of 20 genes involved with “regulation of innate immune response” and “response to a virus” (**Figure 6H**). In addition, the GO terms “defense response to protozoan” and “interferon-mediated signaling pathway” were identified in cultures that received EVs from infected neurons compared to EVs from uninfected neurons and signify the specificity achieved in this stimulation despite no actual parasite being in these cultures (**Figure 6H**). Overall, this data reveals the potential for infected neurons to regulate the inflammatory actions of astrocytes via EVs and to communicate the presence of intracellular Toxoplasma.

## 4 Discussion

Here we demonstrate that extracellular vesicles (EVs) can be harvested from Toxoplasma cyst-containing cortical neuron cultures and that infection of neurons leads to a decrease in EV production and a fundamental change in their content. Significantly, 9 parasite proteins, including GRAs known to be secreted effectors are detected in EVs harvested from cyst-containing neurons. Intact labelled EVs were taken up by astrocytes and observed in the cytoplasm and nucleus of the host cell. Furthermore, incubation of astrocytes with EVs from infected neurons led to a significant change in astrocyte transcriptional signatures, stimulating a pro-inflammatory immune response and downregulating the glutamate transporter GLT-1.

Both our NTA and CD63 ELISA results reveal that infection with *T. gondii* leads to a change in either EV production or release, which results in a significant decrease in the concentration of EVs. All eukaryotic cells release EVs (Schnatz et al., 2021; Silva et al., 2018), therefore infection of neurons could have led to an increase in the concentration of EVs in the media with multiple cells – parasites and neurons – contributing to their production (Augusto et al., 2020; Ben Chaabene et al., 2021; Tomita, Guevara, et al., 2021; Zhang et al., 2022). However, our finding of reduced EV production from cyst-containing neurons is supported by studies from the Carruthers group demonstrating Toxoplasma’s ability to sequester host ESCRT proteins in a GRA14 dependent manner (Rivera-Cuevas et al., 2021; Rivera-Cuevas & Carruthers, 2023). The ESCRT pathway, involved in autophagy and cell division, is also required for extracellular vesicle formation and release (Juan & Fürthauer, 2018). Thus, our finding of reduced EV production from cyst-containing neurons would support the concept that the parasite is reducing the ability of the host cell to operate this machinery by diverting ESCRT proteins for its own use. Although these prior studies showed the highest enrichment of GRA14 bound host proteins in HFF cells, neuronal infection with tachyzoites also led to components of ESCRT I and III binding. Our data would support the ability of Toxoplasma to reduce ESCRT host activity in neurons, even as a slow growing cyst. Alternatively, infection could lead to a reduction in EVs from neurons independent of direct parasite manipulation. Indeed, only half of the neurons in these cultures are infected. Production of EVs can also lead to autocrine uptake (Schnatz et al., 2021). Therefore, an increase in EV uptake by the infected neuron, would also lead to this overall decrease in EV concentration. The mechanism of EV uptake by neurons is unknown but our analysis of protein content of the EVs suggests an increase in “synaptic vesicle endocytosis,” from the upregulation of actin, beta (Actb) and clathrin heavy chain (Cltc), (**Figure 3D**) and many proteins involved in the uptake of synaptic vesicles are also involved in the numerous potential mechanisms of EV uptake (Mulcahy et al., 2014).

The third potential reason for a decrease in EV’s harvested from infected neurons is that infection is causing a change in neuronal health and function. Indeed, this occurs *in vivo* as reported by several groups (Correa Leite et al., 2021; David et al., 2016; Mendez et al., 2021; Tedford & McConkey, 2017). These short-term *in vitro* cultures, long enough for cyst formation, do not lead to the death of neurons however a reduction in dendrite formation can be seen by immunocytochemistry (**Figure 1A**; **Supplemental Figure 1A/C**). In addition, our data revealed the downregulation of miR-324-5p in EVs following infection (**Figure 4**; **Supplemental Table 3**), which when knocked out in mice can lead to a decrease in dendritic spine density and morphology in the brain (Parkins et al., 2023, 2023). Thus, infected neurons are retracting dendrites, becoming less morphologically complex and such a process, involving a reduction in surface area and membrane modifications may also be associated with reduced EV release.

There is an assumption that the measurement of proteins and small molecules in EVs directly and proportionally reflects the content and function of the cell with proteins being indiscriminately packaged into the vesicle. However, there is evidence of selectivity and sorting of cargo prior to EV release, for example the specific exclusion of pro-inflammatory mitochondrial proteins (Anand et al., 2019; Todkar et al., 2021). In addition, the purported function of EVs as an important mechanism of cell-cell communication suggests a need for packaging of cargo to be targeted and selective.

Our investigation of the content of EVs from infected neurons revealed significantly altered protein and miRNA content. Heat-shock proteins are common stress response regulators and are also protein markers of EVs. Our data reveals their presence and an increase in HSP90 and Hsp90b1 following infection (De Maio & Vazquez, 2013; Saito et al., 2024; Sojka et al., 2023). In addition, we are further able to confirm that these are neuronal EVs through the presence of neural cell adhesion molecule L1-like protein (Chl1) (Solana-Balaguer et al., 2023). While many of the proteins were not modified upon infections, there were several proteins that were significantly up or downregulated. Overall the changes in EV protein content following Toxoplasma infection suggests a downregulation of neuronal organization and migration, picked up by two downregulated GO terms and includes the 4-fold reduction in reelin (Reln), a glycoprotein particularly important for neuronal migration during development with its decrease often an indicator of developmental disorders (Alcántara et al., 2006; Wierman et al., 2011). Morphological alterations and a decrease in neuronal migration is consistent with previously published work (Pires et al., 2023) and is supported by the appearance of neurons in culture (**Figure 1**) and the contraction and decrease in neuronal complexity *in vivo* (David et al., 2016; Mendez et al., 2021).

Infection led to EV content that would suppress a pro-inflammatory status. MicroRNAs are frequently inhibitory but can also be activators of gene transcription (Odame et al., 2021). The miRNA content of EVs from infected neurons is overwhelmingly associated with blocking NF-κB and other pro-inflammatory pathways. Both the downregulation of miR-3473b, (X. Wang et al., 2018) and upregulation of miR-199a-3p, miR-143-3p and miR-21a-5p (Duan et al., 2022; Peng et al., 2020; Y. Wang et al., 2020; Xiong et al., 2019; Ye et al., 2018) leads to a reduction in inflammatory signals. Notably, miR-199a-3p, upregulated over 5-fold in EVs from infected neurons, has been reported to suppress astrocyte activation and NF-κB-mediated inflammation (Saadh et al., 2023), while miR-143-3p negatively regulates NOTCH signaling and in this context may, as a result, decrease cell proliferation, differentiation and BBB integrity (Bai et al., 2016; Hu et al., 2023). This suggests that miRNA-143-3p is benefiting the infection and spread of the parasite in the brain.

In addition to proteins, miRNA-143-3p and miRNA-21a-5p have been upregulated in Toxoplasma infected macrophages and microglia EVs, and have been predicted to regulate nitric oxide (NO) synthase (Cong et al., 2017; Jung et al., 2022; S. Li et al., 2019). In addition, miR-424-5p upregulation during cerebral injury has led to neuroinflammation and apoptosis (Luo et al., 2022), while miR-30d-5p also increases apoptosis and autophagy during brain ischemia (Zhao et al., 2017). Another protein only identified in EVs from infected neurons and highly upregulated is vimentin (Vim). Vim is an intermediate filament protein found in the cytoskeleton that is able to inhibit type I interferon production (Liu et al., 2022), and in mouse models of spinal cord injury, Vim has also been identified as a key regulator in reactive astrocytes (Q. Li et al., 2024). There is a significant increase in vimentin in EVs following *T. gondii* infection, indicating that these EVs may promote an anti-inflammatory immune response.

By using the resources available on ToxoDB (https://toxodb.org) we were also able to determine one of the most revealing aspects of EV content analysis namely the presence of parasite specific proteins. A total of 8 parasite proteins were identified in EVs from infected neurons. Three of these, HSP90, TUBA1 and GAPDH1, are found in EV’s from many eukaryotic cells. However, an additional 5 proteins, GRA1, 2, 7 and MAG1 and 2 play a role in cyst formation but importantly are secreted by the parasite and many have the potential to manipulate the host immune response. GRA proteins help form the intravacuolar network (IVN), cyst wall and the inner cyst matrix (Guevara et al., 2019). Our EVs are harvested after 3 days of cyst formation and therefore these cysts are relatively immature. GRA1, found at one of the highest levels in EVs, at this time point is found on the outer layer of the cyst, is not pulled down in the presence of CST1 (carbohydrate components of the tissue cyst) and is dispensable for cyst growth and stability (Guevara et al., 2019; Tu et al., 2019). This and the presence of GRA1 in EVs could support the concept that GRA1 is a protein that is secreted by bradyzoites and interacts with the host cell rather than playing a critical structural or developmental role. GRA2 as well as being at the periphery is found preferentially in the cyst wall matrix and remains there throughout the maturation of the cyst (Guevara et al., 2019). Both GRA1 and GRA2 are associated with increased cyst survival in the presence of T cells (Mani et al., 2023) supporting a role for host cell manipulation and suppression of immunity. Indeed, GRA7, the third most abundant GRA in EVs from infected neurons, is a protein known to interact with the host, and when phosphorylated interacts with GRA1 (Dunn et al., 2008; Yang et al., 2016). In addition, MAG1 and MAG2, both components of the cyst matrix, are seen by the immune system as evidenced by the significant anti-MAG immune response (Carruthers & Suzuki, 2007; Tu et al., 2020; Yang et al., 2016). MAG1, the most abundant parasite derived protein in our EVs is secreted and can suppress inflammasome activation (Tomita, Mukhopadhyay, et al., 2021). Importantly, similar proteins seen during cyst development (e.g. GRA12) were not found in our EVs suggesting either that these do not escape the cyst or that there is selectivity in the parasite-derived cargo of EVs. Thus, the parasite proteins found in infected neuron-derived EVs are known secreted proteins with many of them capable of interacting with the host cell.

Our original goal was to determine whether infection of neurons could indirectly lead to the downregulation of GLT-1 in astrocytes via changes in neuronal EVs (David et al., 2016; Men et al., 2019; Morel et al., 2013). Although there was no change in miRNA-124 in the EVs following infection, indicating that GLT-1 regulation is at least partially miRNA-124 independent, we did demonstrate that the addition of EVs from infected neurons significantly decreases GLT-1 protein expression in primary astrocytes. Our data demonstrate that labelled EVs are taken up by astrocytes in culture and colocalize not only to the cytoplasm but also the nucleus of the cell (**Figure 5**). Other studies have reported observing EVs within the nucleus with incorporation into late endosomes being the likely mechanism (Corbeil et al., 2020; Rappa et al., 2017). Although we have ruled out that initial uptake of neuronal EVs by astrocytes is via macropinocytosis (Fitzner et al., 2011; Pantazopoulou et al., 2023), further studies will need to be conducted to determine the uptake mechanism. Despite the content of EVs suggesting their potential to limit inflammation or be ‘pro-parasite’, addition of EVs from infected neurons altered astrocyte gene expression towards a pro-inflammatory immune response. GO analysis revealed a large number of genes involved in immune defense and interferon mediated signaling. Astrocytes are able to control parasite replication through the activation of STAT1 (signal transducer and activator of transcription 1) and the use of IRGs (immunity regulated GTPases) (Halonen et al., 2001; Rosowski et al., 2014) and are thus very adept at responding to Toxoplasma. It implies that the EV parasite-derived cargo of GRA proteins, instead of manipulating the host cell are being detected in a classical immune mediated mechanism either by pattern recognition receptors or inflammasome activation. Alternatively, the combined changes in host neuronal proteins and nucleic acids in the EVs could be indicating that the source cell requires an astrocytic change in function from a supportive neurotransmitter regulating cell to a damage control agent.

This study tested the effect of neuronal EVs on astrocytes due to the known close relationship between these cells and the specific questions on the infection-induced changes in GLT-1. However, EVs derived from cyst containing neurons could also be taken up by many other cells in the brain including microglia and peripheral immune cells and therefore this alteration in host EV production and content in a few infected neurons could have a much broader impact on neuroinflammation in the brain.

Lastly, the use of EVs as known biomarkers for disease, especially as readouts of what is taking place in the brain, is a growing possibility (Xu et al., 2024). EVs generated in the CNS can be detected in cerebral spinal fluid, blood, tears, and urine. As we have seen in our own data, they provide cell specificity with neuronal derived EVs containing Chl1. Toxoplasma infection is determined by the presence of peripheral circulating antibodies that would be generated no matter the location of infection. A large body of work has attempted to correlate seropositivity with other unrelated neurological conditions despite not knowing whether infection includes cysts in the brain (Savitz & Yolken, 2023). The demonstration that EVs from infected neurons contain stage-specific parasite proteins provides a potential screen for the presence of neuronal cysts and a refinement of the broader effects of Toxoplasma infection on brain neurochemistry and clinical disease.

Overall, our data suggests that the presence of Toxoplasma cysts within neurons has the potential to broadly change the brain environment through an alteration in the production and content of extracellular vesicles. The dispersion and uptake of these EVs drives an infection-induced astrocyte phenotype promoting inflammatory signaling at the expense of neurotransmitter uptake.

## Author Contribution

**Emily Z. Tabaie:** Conceptualization; Data curation; Formal analysis; Investigation; Methodology; Visualization; Validation; Writing - original draft, review & editing. **Ziting Gao:** Methodology; Resources. **Stacey Gomez:** Data curation; Methodology. **Kristina Bergersen**: Data curation; Methodology. **Wenwan Zhong:** Supervision; Resources. **Emma H. Wilson:** Conceptualization; Funding acquisition; Project administration; Supervision; Writing - review & editing.

## Acknowledgements

This work was supported, in part, by funds from the Division of Biomedical Sciences at the University of California, Riverside. Bioanalyzer, Library preparation, and sequencing services on miRNA provided by the Genomics Core, Institute for Integrative Genome Biology, UC Riverside. Matthew Dickson at the University of California, Riverside, Advanced Microscopy and Microanalysis core is acknowledged for performing TEM on the EVs. Brandon Le at the University of California, Riverside, Genomics Core for assistance with the RNASeq data analysis. Quanqing Zhang at the University of California, Riverside, Proteomics Core for running liquid chromatography/mass spectrometry on the EVs. This publication includes data generated at the UC San Diego IGM Genomics Center utilizing an Illumina X Plus that was purchased with funding from a National Institutes of Health SIG grant (#S10 OD026929) and the use of the High-Performance Computing Center (HPCC) at UC Riverside (NSF#2215705). All authors would like to thank the fundamental resource that is ToxoDB and hope for its continuation.

## Supporting Information

Additional supporting information can be found online in the Supporting Information section at the end of this article.

## Data Availability

The authors declare that all data supporting the findings of this study are available within the article and its supplementary information files. Raw sequencing and processing files for RNA and miRNA Sequencing have been deposited in the GEO database under accession code: PRJNA1158965. Raw and processed files for LC-MS/MS protein analysis have been deposited in the MassIVE database under accession code: MSV000096222.

